# An event-related potential study of decision making and feedback utilization in female college students who binge drink

**DOI:** 10.1101/582734

**Authors:** Eun-Chan Na, Kyoung-Mi Jang, Myung-Sun Kim

## Abstract

This study investigated the ability to use feedback for decision making in female college students who binge drink (BD) using the Iowa Gambling Task (IGT) and event-related potentials (ERPs). Twenty-seven binge drinkers and 23 non-binge drinkers (non-BD) were identified based on scores on the Korean version of the Alcohol Use Disorder Test and the Alcohol Use Questionnaire. The IGT consists of four cards, including two cards that result in a net loss, with large immediate gains but greater losses in the long term, and two cards that result in a net gain, with small immediate gains but reduced losses in the long term. Participants were required to choose one card at a time to maximize profit until the end of the task while avoiding losses. The BD group showed a significantly lower total net score than the non-BD group, indicating that the BD group chose more disadvantageous cards. The BD group showed significantly smaller ΔFRN amplitudes (difference in amplitudes of feedback-related negativity [FRN] between gain and loss feedback) except in P3. Additionally, ΔFRN amplitudes in the fronto-central area were positively correlated with the total net score and net scores for sectors 4 and 5. Thus, total net scores and later performance on the IGT increased as ΔFRN amplitudes from the fronto-central area increased. FRN is known to reflect early feedback evaluation employing a bottom-up mechanism, whereas P3 is known to reflect late feedback processing and allocation of attentional resources using a top-down mechanism. These results indicate that college students who binge drink have deficits in early evaluation of positive or negative feedback and that this deficit may be related to decision making deficits.

## Introduction

Binge drinking (BD) is defined as a repeated pattern of excessive alcohol consumption and abstinence over a short period of time [1–3]. BD is most prevalent among young adults, especially college students [3,4,5], and is associated with various problems including assault, drunk driving, unguided or unsafe sexual behavior, and academic underachievement [3,6–8]. Additionally, binge drinkers exhibit similar structural and functional brain abnormalities and neuropsychological deficits to patients with alcohol use disorder (AUD) [1,9–13], and BD predicts the development of AUD in the future [11,14–16].

Patients with AUD cannot stop drinking alcohol even though they suffer from its negative consequences [17,18]. Such behaviors reflect inefficient decision making among patients with AUD, as they continue to seek immediate rewards and ignore future consequences [19,20]. In other words, they not only underestimate the negative consequences of alcohol consumption [21] but also emphasize immediate rewards over long-term consequences [22,23]. Decision making deficits have been observed in patients with AUD [19,24–28] and in binge drinkers [29–33].

Decision making is defined as a process of forming a preference for an option, making a choice based on the preference, executing the choice, and evaluating the consequences of the choice [34]. Decision making is a complex process including both cognitive and non-cognitive processes (i.e., emotions) [35], and various brain areas, such as the orbitofrontal, ventromedial prefrontal, anterior cingulate cortices, and amygdala, are involved in decision making [36–40].

The Iowa Gambling Task (IGT) is widely used to evaluate decision making ability [41,42]. Participants are asked to choose one of four cards on every trial to maximize profit while avoiding loss. The chosen card results in gains on every trial, but also results in intermittent losses. The cards differ in feedback magnitude and probability. Two cards (A and B) result in large immediate gains but greater losses, causing a net loss (disadvantageous cards), whereas the other two cards (C and D) lead to small immediate gains and smaller losses, resulting in a net gain (advantageous cards). Participants must evaluate feedback such as valence (gain or loss), magnitude (large or small), and the probability of encountering losses to learn the contingency between the card and its consequences [43,44].

Studies investigating decision making ability in patients with AUD using the IGT found that patients with AUD performed poorly compared to normal controls, choosing significantly more disadvantageous cards and significantly fewer advantageous cards compared with the controls [19, 25,26,28,45]. Additionally, positive correlations were observed between IGT performance and grey matter volume in the dorsal and ventromedial prefrontal cortices, which are crucial for decision making [46]. Poor IGT performance has also been observed in individuals with BD [29,30,33,47]. For example, adolescents [47] and college students with BD [33] performed significantly worse on the IGT than did non-BD group.

Feedback utilization, a process of identifying whether an action induces positive or negative consequences and evaluating those consequences, is crucial to making efficient decisions [48]. Considerable improvement in our understanding of the neurological basis of feedback utilization has revealed that the orbitofrontal, ventromedial prefrontal, and anterior cingulate cortices as well as the ventral striatum are involved in feedback utilization [49–53]. The ventral striatum is involved in prediction errors, i.e., how actual feedback differs from personal expectations, whereas the orbitofrontal cortex is involved in evaluating feedback based on prediction errors [54–57]. Additionally, the anterior cingulate cortex evaluates rewards in situations where contingencies are uncertain and then relays the evaluation of the reward to motor areas for response execution [37].

Studies using event-related potentials (ERPs) suggest two components, feedback-related negativity and P3, as the electrophysiological indices of feedback utilization [48,58]. Gehring and Willoughby [58] used a simple gambling task to observe a negative peak approximately 265 ms post feedback whose amplitude was larger in response to negative than to positive feedback. This peak is known as feedback-related negativity (FRN) or outcome-related negativity [59]. FRN is sensitive to feedback valence (gain or loss) [60] and is associated with activation of the midbrain dopaminergic system [61]. Additionally, reinforcement-learning theory suggests that FRN reflects prediction errors, i.e., the difference between actual feedback and personal expectation [48,62,63]

P3, another ERP component related to feedback utilization, is a positive peak observed in central-parietal areas at 275-700 ms post feedback [48,59]. P3 is known to be sensitive not only to feedback valence but also to feedback magnitude and probability [60,63–66]. It has been suggested that P3 reflects activation of the locus coeruleus-norepinephrine system and processing of task-relevant information to maximize decision making efficiency [67]. In other words, P3 reflects, unlike FRN, a top-down mechanism that processes and evaluates feedback-related information in detail [48,63].

Alcohol consumption affects feedback utilization. A study that used a gambling task and measured ERPs found that the alcohol consumption group exhibited significantly lower FRN amplitudes in response to both gain and loss feedback, especially to loss feedback, than did a placebo group, indicating that alcohol consumption affects feedback utilization [68]. Deficits in feedback utilization are also observed in patients with AUD. For example, Fein and Chang [69] using the Balloon Analogue Risk Task, observed that patients with AUD and a family history of AUD exhibited significantly smaller FRN amplitudes than did those without a family history. Kamarajan et al. [70] used a gambling task and reported that patients with AUD exhibited lower P3 amplitudes in response to both gain and loss feedback and smaller FRN amplitudes to loss feedback than did normal controls. Additionally, they observed increased activation in primary sensory and motor areas during the FRN time window and decreased activation in the cingulate gyrus during the P3 window in patients with AUD relative to normal controls. These results indicate that the sensory and motor areas of patients with AUD are hyper-excited during early feedback evaluation, and areas involved in feedback evaluation are hypo-activated compared to normal controls [70].

To our knowledge, only one study has investigated feedback utilization deficits in binge drinkers using ERPs. That study, which used the IGT, found that the BD group tended to exhibit smaller FRN amplitudes (*p* = .06) than the non-BD group [71]. However, that study used the original computerized IGT [41,42], which had two limitations: First, the original IGT consisted of 100 trials, which is not suitable for an ERP study where a sufficient number of trials is needed [72]. Second, the original IGT displays gains in every trial and subsequently displays losses according to each card’s probability. When multiple stimuli are displayed in succession, the ERPs to loss feedback might be contaminated by previous gain feedback.

The present study investigated feedback utilization ability during decision making in BD female college students using the IGT and ERP. Specifically, this study examined whether decision making deficits in BD female students are related to feedback utilization deficits and, if so, how they are reflected in feedback-related ERP components, FRN and P3. Based on previous findings, we hypothesized that the BD group would perform significantly worse than the non-BD group on the IGT; that the BD group would show significantly smaller FRN and P3 amplitudes than the non-BD group; and that IGT performance and feedback-related ERPs would be positively correlated. As gender differences are observed in BD [73–75], decision making [76], and ERP amplitudes [77], only female college students were included in this study.

## Materials and methods

### Participants

The details of the participant screening procedures have been described in previous studies by our research group [33,78]. The Korean version of the Alcohol Use Disorder Identification Test (AUDIT-K) [79,80], Alcohol Use Questionnaire (AUQ) [81], and a questionnaire inquiring about binge drinking episodes in the last 2 weeks were administered to 435 female college students. The BD and non-BD groups were defined based on 1) alcohol-related problems and drinking habits, 2) the number of BD episodes, and 3) drinking speed. The BD group included those who 1) scored at least 12 but less than 26 on the AUDIT-K, 2) had consumed four or more glasses at one sitting in the last 2 weeks, and 3) drank two or more glasses per hour. Although the World Health Organization (WHO) recommends using a score > 8 as the cutoff point for problem drinking [79], the cutoff score of 12 was applied because a cutoff point of 8 includes those who do not have apparent drinking problems but may display problem drinking in the future [82,83]. In contrast, those who received scores > 26 on the AUDIT-K were also excluded, as AUD was suspected. The non-BD group included those who 1) scored less than 8 on the AUDIT-K, 2) had not drunk four or more glasses in one sitting in the last 2 weeks, and 3) drank 1 glass or less per hour.

The Structured Clinical Interview of the Diagnostic and Statistical Manual of Mental Disorders-Fourth Edition (SCID-NP) [84] was administered to ensure that no participants had a psychiatric disorder. Additionally, the Self-rating Depression Scale (SDS) [85], the State-Trait Anxiety Inventory (STAI) [86], and Barratt Impulsivity Scale (BIS) [87] were administered to evaluate depression, anxiety, and impulsivity, respectively. To control for the influence of alcohol-related genes and family history, the Korean version of the Children of Alcoholics Screening Test (CAST-K) [88,89] was administered, and those who scored 6 or more were excluded. Last, those who were left-handed or ambidextrous were also excluded to control for the effect of brain lateralization.

In the end, 50 students participated in this study (27 in the BD group and 23 in the non-BD group). This study was approved by Sungshin Women’s University Institutional Review Board (SSWUIRB 2017-040). The participants provided written informed consent after receiving a description of the study, and they were paid for their participation.

### The Korean version of the Alcohol Use Disorder Identification Test (AUDIT-K)

The AUDIT [79], a self-administered questionnaire designed to measure the presence of AUD and drinking problems, consists of 10 items. The total score ranges from 0 to 40. Three items inquire about frequency and quantity of alcohol consumption, three about symptoms related to alcohol dependence, and four about psychosocial problems related to alcohol consumption. The Korean version was administered in this study [80].

### Alcohol Use Questionnaire (AUQ)

The AUQ [81] is a self-administered questionnaire measuring dinking patterns. Items 10, 11, and 12 evaluate drinking speed, frequency of being drunk within the last 6 months, and the rate of being drunk when consuming alcohol, respectively. These three items were used to calculate a BD score [90]. The binge score was calculated using the following equation:

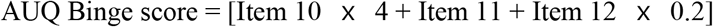

### The Iowa Gambling Task (IGT)

This study employed a modified version of the original computerized IGT [41] to make the task suitable for measuring ERPs (Fig 1A). Four cards were displayed on a computer monitor, and participants were asked to maximize profits until the end of the game by choosing a card during each trial. Gain or loss feedback was displayed after each choice, with gain feedback consisting of a green smiling emoticon with points earned and loss feedback consisting of a red crying emoticon with the points lost (Fig 1B).

**Fig 1.**
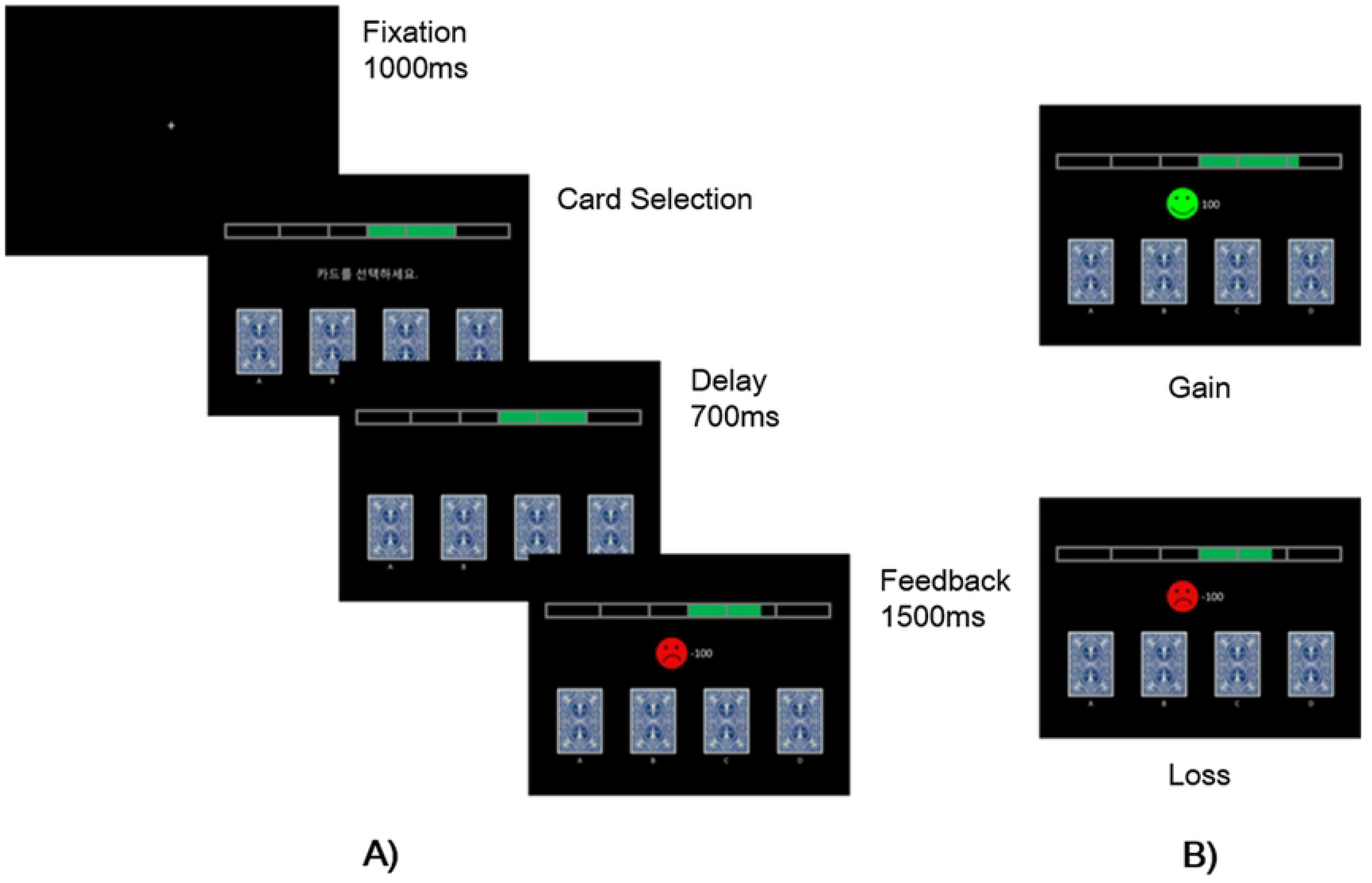
The modified IGT. A) A fixation point will be displayed for 1,000ms then four cards will be displayed till the participants make their choice. At 700ms after a card is chosen, feedbacks will be displayed for 1,000ms. B) The feedback stimuli consists of gain conditions and loss conditions. In gain conditions, green smiling emoticon and the earned points will be displayed whereas red crying emoticon and the lost points will be displayed in loss conditions.

The magnitude and probability of gain and loss for each card were set as for the original computerized IGT [41]. The cards consisted of two disadvantageous cards (A and B), which provided large gains and larger losses, resulting in a net loss, and two advantageous cards (C and D), which provided small gains but smaller losses, resulting in a net gain. Cards A and C each had a 50% chance of causing losses, whereas cards B and D had a 10% chance of causing losses.

The task consisted of three blocks; the locations of the cards were changed at the beginning of each block to keep participants motivated. Each block comprised 100 trials; a total of 320 trials, including 20 practice trials, were administered. Decision making ability was measured by the net score, which was calculated by subtracting the frequency of choosing the disadvantageous cards (A and B) from the frequency of choosing the advantageous ones (C and D)

E-Prime software (version 2.0; Psychological Software Tools, Inc., Sharpsburg, PA, USA) was used to administer the modified IGT. A fixation point (+) was displayed for 1,000 ms, and the cards were then displayed until the participants made their choice by pressing a button. The feedback, either a gain or loss, was displayed for 1,000 ms at 700ms after a card was chosen.

### Electrophysiological recording procedure

Electroencephalography (EEG) was measured using a 64-channel Geodesic sensor net connected to a 64-channel, high-input impedance amplifier (Net Amp 300; Electrical Geodesics, Eugene, OR, USA) in a shielded and soundproofed room. All electrodes were referenced to Cz, and impedance was maintained at 50 KΩ or less [91]. EEG activity was recorded continuously using a 0.3 - 100 Hz bandpass filter at a sampling rate of 500 Hz. The recorded EEG data were digitally filtered using a 0.3 - 30 Hz bandpass and re-referenced to the average reference. The continuous EEG was then segmented into 800 ms epochs (from 100 ms pre-to 700 ms post-feedback). Additionally, epochs contaminated by artifacts such as eye blinks were removed based on the threshold of a peak-to-peak amplitude of ± 70 μV from the eye channels. The remaining data were averaged according to feedback valence, i.e., gain and loss feedback.

### Statistical analysis

Demographic variables were analyzed with independent *t*-tests. The total net scores on the modified IGT were analyzed with independent *t*-tests. Additionally, each block was subdivided into five sectors, and scores for each sector were averaged across the three blocks to calculate sector net scores to measure performance improvement across trials. The sector net scores were analyzed with mixed-design analysis of variance (ANOVA), where group (BD or non-BD) was a between-subjects factor, and sector (1 - 5) was a within-subject factor.

ERP components and time windows were determined based on grand averaged ERPs and individual ERP waveforms. FRN was defined as the most negative peak observed at 200 - 275 ms after feedback-onset, and P3 was defined as the most positive peak followed by FRN, i.e., observed 275 - 600 ms after feedback. Because the FRN and P3 time windows overlapped and because the FRN is a negative and P3 is a positive peak, it is possible that latent components representing FRN and P3 independently might be distorted on the ERP waveforms due to the overlapping windows where the amplitudes and latencies do not clearly represent the differences by feedback valence [92]. To overcome this problem, it is necessary to isolate ERP components; difference waves have been recommended for this purpose [92]. Therefore, ∆FRN (FRN effect) and ∆P3 (P3 effect) were defined as the amplitude difference between gain and loss feedback [64,66,93–96].

Amplitudes and latencies of each component were analyzed by mixed ANOVA. Electrode site (FC3, FCz, FC4, C3, Cz, C4, P3, Pz, and P4) and valence (gain or loss) were within-subject factors, and group was a between-subjects factor. The electrode sites for ∆FRN and ∆P3 were a within-subject factor, and group was a between-subjects factor. Greenhouse-Geisser corrections were used in cases of violation of sphericity, and corrected *p*-values are reported when appropriate. The mean numbers of trials included in the FRN/P3 analysis for the BD and non-BD groups were 105.57 (gain = 161.89, loss = 51.26) and 111.33 (gain = 17.48, loss = 52.17), respectively. The two groups did not differ in terms of trials for averaging FRN/P3 in the gain feedback (F[1,48] = .62, *p* = .44), the loss feedback (F[1,48] = .04, *p* = .85) or both feedbacks (F[1,48] = .57, *p* = .46). The relationships of the ∆FRN and ∆P3 amplitudes with performance on the IGT, i.e., total net scores and sector net scores, were analyzed using Pearson’s correlation coefficient analysis. A *p*-value < .05 was considered significant.

## Results

### Demographic characteristics

The BD and non-BD groups did not differ in terms of age (t[48] = −1.08, *p* = .29), educational level (t[48] = −1.07, *p* = .29), SDS (t[48] = .80, *p* = .43), or trait anxiety on the STAI (t[48] = 1.05, *p* = .30). However, the BD group exhibited significantly higher state anxiety on the STAI (t[48] = 5.49, *p* < .001), BIS (t[48] = 6.92, *p* < .001), AUDIT-K total score (t[48] = 16.81, *p* < .001), drinking speed (t[48] = 12.56, *p* < .001), frequency of being drunk within the last 6 months (t[48] = 5.63, *p* < .001), percentage of being drunk when consuming alcohol (t[48] = 3.73, *p* < .01), and AUQ binge score (t[48] = 9.94, *p* < .001) compared to the non-BD group. The demographic characteristics of the BD and non-BD groups are presented in Table 1.

**Table 1.** Demographic characteristics of the non-BD and BD groups.

| | Non-BD<br>($n = 23$) | BD<br>($n = 27$) | $t$ |
| --- | --- | --- | --- |
|  | Mean (SD) | Mean(SD) |  |
| Age (years) | 22.04(1.92) | 21.44(1.99) | -1.08 |
| Education (years) | 15.09(1.08) | 14.74(1.20) | -1.07 |
| SDS | 39.61(5.39) | 41.15(7.83) | .80*** |
| STAI state | 38.57(8.10) | 56.93(14.16) | 5.49** |
| STAI trait | 38.70(7.56) | 41.26(9.37) | 1.05 |
| BIS | 63.48(10.95) | 83.26(9.29) | 6.92** |
| AUDIT-K | 2.39(1.80) | 17.37(4.20) | 16.81*** |
| Speed of drinking<br>(drinks/hour) | .65(.57) | 4.22(1.34) | 12.56*** |
| Times drunk in the last<br>6 months | .13(.34) | 5.07(4.55) | 5.63*** |
| Percentage of times<br>became drunk when<br>drinking (%) | 11.87(23.37) | 39.44(28.83) | 3.73** |
| AUQ binge drinking<br>score | 5.11(5.17) | 29.85(11.66) | 9.94*** |

**Notes:** \*\* $p < .01$ \*\*\* $p < .001$ .
**Abbreviations:** SDS, Self-Rating Depression Scale; STAI, Spielberger's State-Trait Anxiety Inventory; BIS, Barratt Impulsivity Scale; AUDIT-K, The Korean version of Alcohol Use Disorder Identify Test; AUQ, Alcohol Use Questionnaire.

As significant differences in state anxiety and impulsivity were detected, mixed analysis of covariance was performed with state anxiety and impulsivity as covariates to control their effect on the IGT and ERP components. However, the analysis revealed that state anxiety as a covariate was not significantly associated with the IGT (*p* = .086), FRN (*p* = .565), or P3 (*p* = .634) and that impulsivity as a covariate was not significantly associated with the IGT (*p* = .464), FRN (*p* = .295), or P3 (*p* = .631).

### The modified Iowa Gambling Task (IGT)

The BD group exhibited a significantly lower total net score than the non-BD group (t[48] = −2.61, *p* < .05). In terms of sector net scores, a main effect of sector was observed (F[4,192] = 2.45, *p* < .05). A further post hoc analysis revealed a trend toward a lower net score for sector 2 than for sector 4 (*p* = .09). Additionally, a main effect of group was observed (F[1,48] = 7.28, *p* < .05), with the BD group exhibiting significantly lower sector net scores than the non-BD group. However, the sector ⅹ group interaction was not significant (F[4,192] = 1.226, *p* = .30). Mean total and sector net scores of the BD and non-BD groups are presented in Table 2 and Fig 2.

**Table 2.** Performance of the modified IGT in the non-BD and BD groups.

| | Non-BD<br>($n=23$) | BD<br>($n=27$) |
| --- | --- | --- |
|  | Mean (SD) | Mean (SD) |
| Sector 1 | -.03 (4.16) | -1.73 (3.04) |
| Sector 2 | .72 (5.73) | -3.04 (4.14) |
| Sector 3 | .93 (5.53) | -1.63 (3.49) |
| Sector 4 | 2.23 (5.75) | -1.41 (4.35) |
| Sector 5 | 1.25 (5.14) | -1.93 (4.08) |
| Total | 5.10 (23.43) | -9.73 (15.11) |

**Fig 2.**
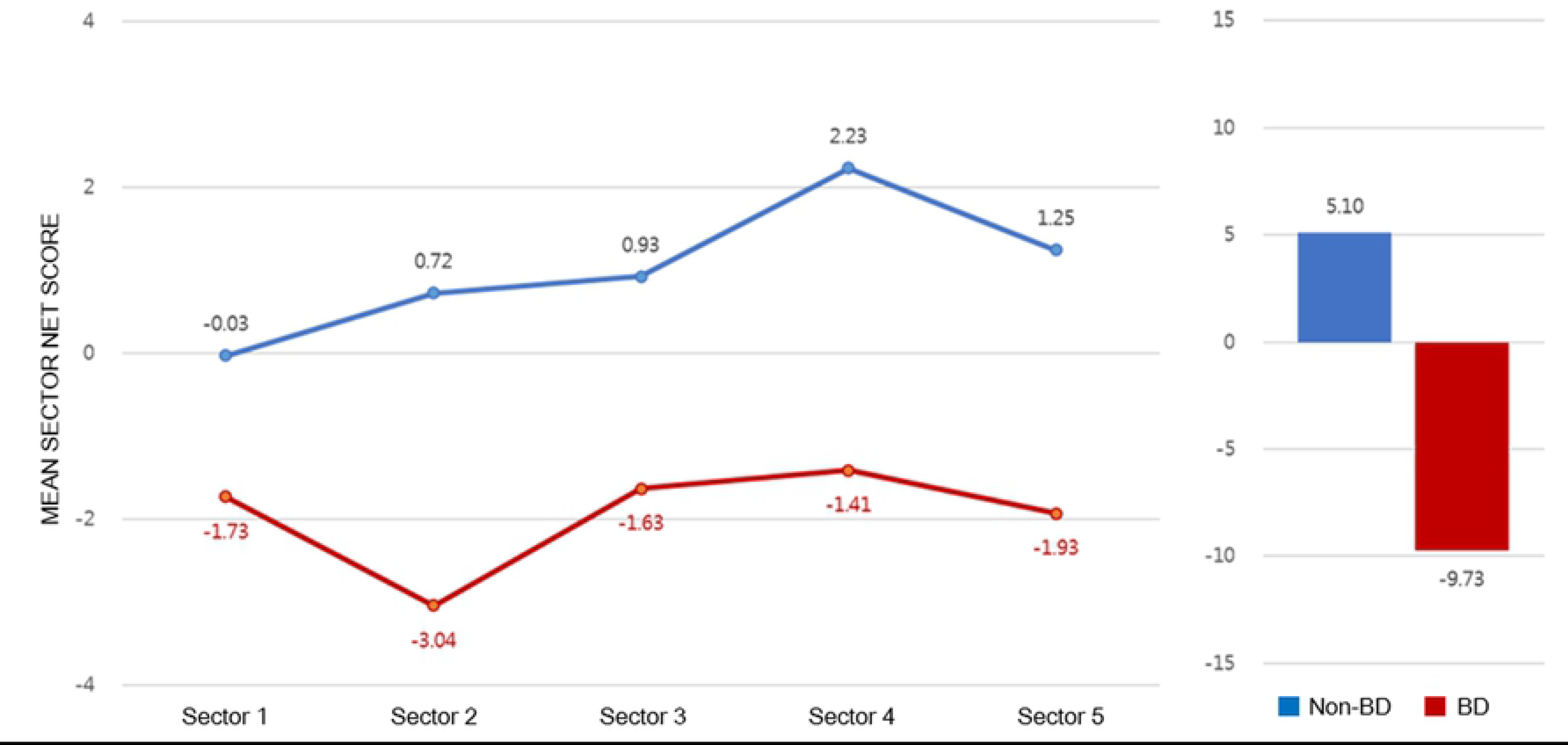
Performance of the modified IGT. Sector net scores (left) and total net scores (right) of the modified IGT in non-binge drinking and binge drinking groups.

### Electrophysiological measures

The grand-averaged ERPs elicited by gain and loss feedback at fronto-central (FCz), central (Cz), and parietal midlines (Pz) for the BD and non-BD groups are displayed in Fig 3. The BD and non-BD groups exhibited the largest FRN and P3 amplitudes at Cz. The topographical distribution of FRN and P3 measured at all electrodes when the largest FRN and P3 amplitudes were observed are displayed in Figs 4 and 5, respectively.

**Fig 3.**
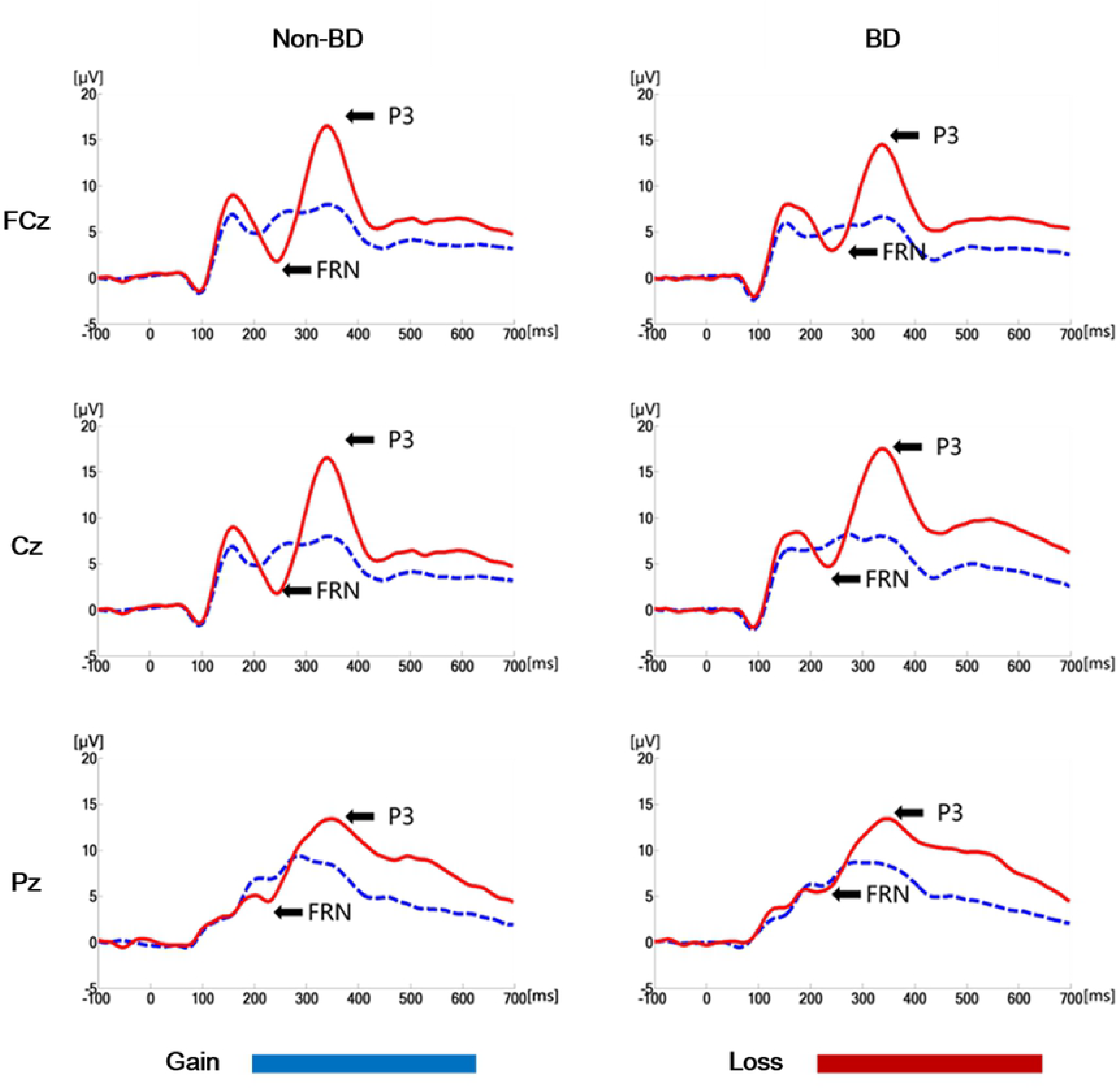
The grand-averaged ERPs. The grand-averaged ERPs elicited by gain and loss feedback at FCz, Cz and Pz for non-binge drinking and binge drinking groups.

**Fig 4.**
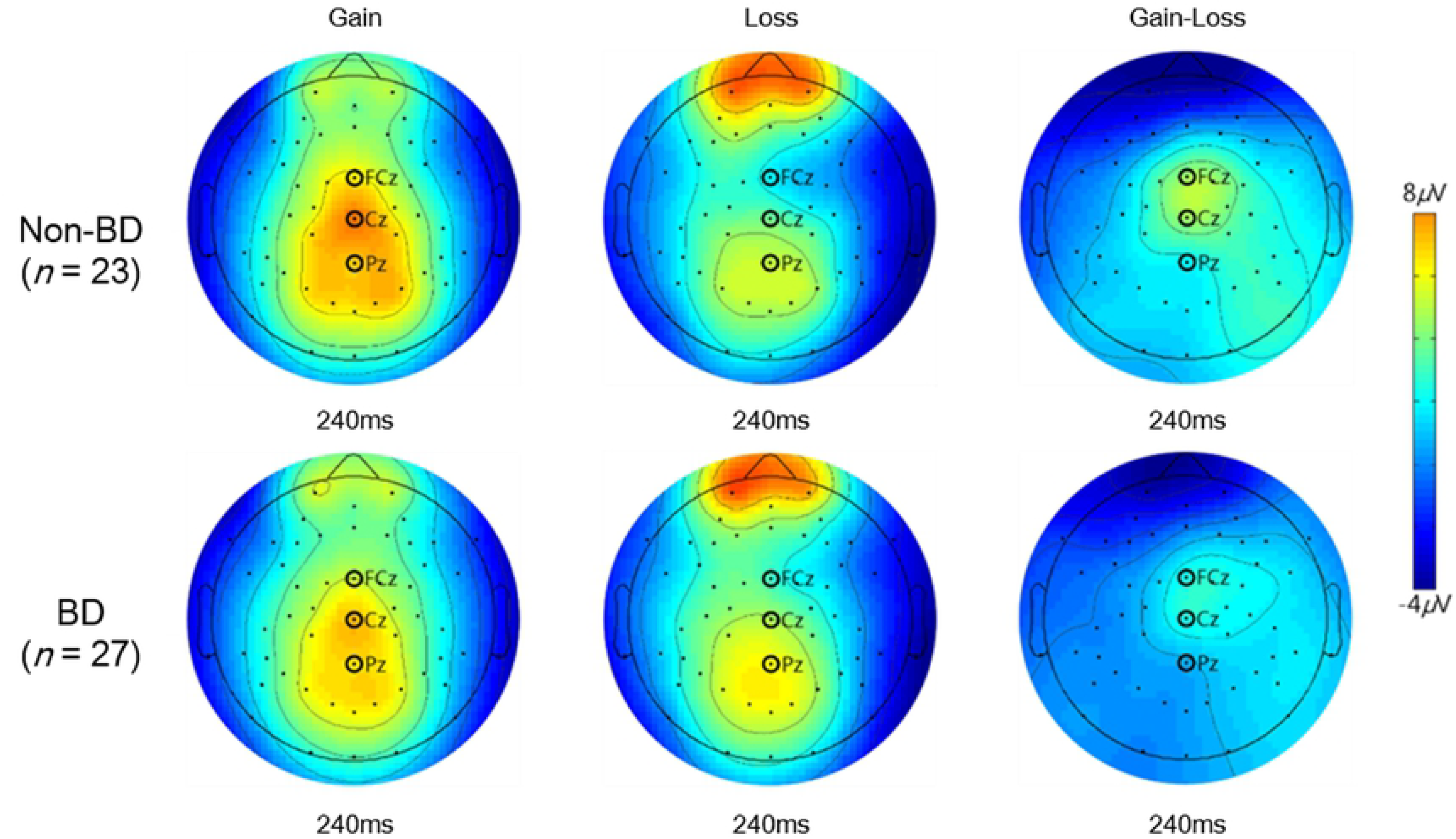
Topographical distribution of FRN. The topographical distribution of FRN measured at all electrodes when the maximum FRN amplitudes were observed.

**Fig 5.**
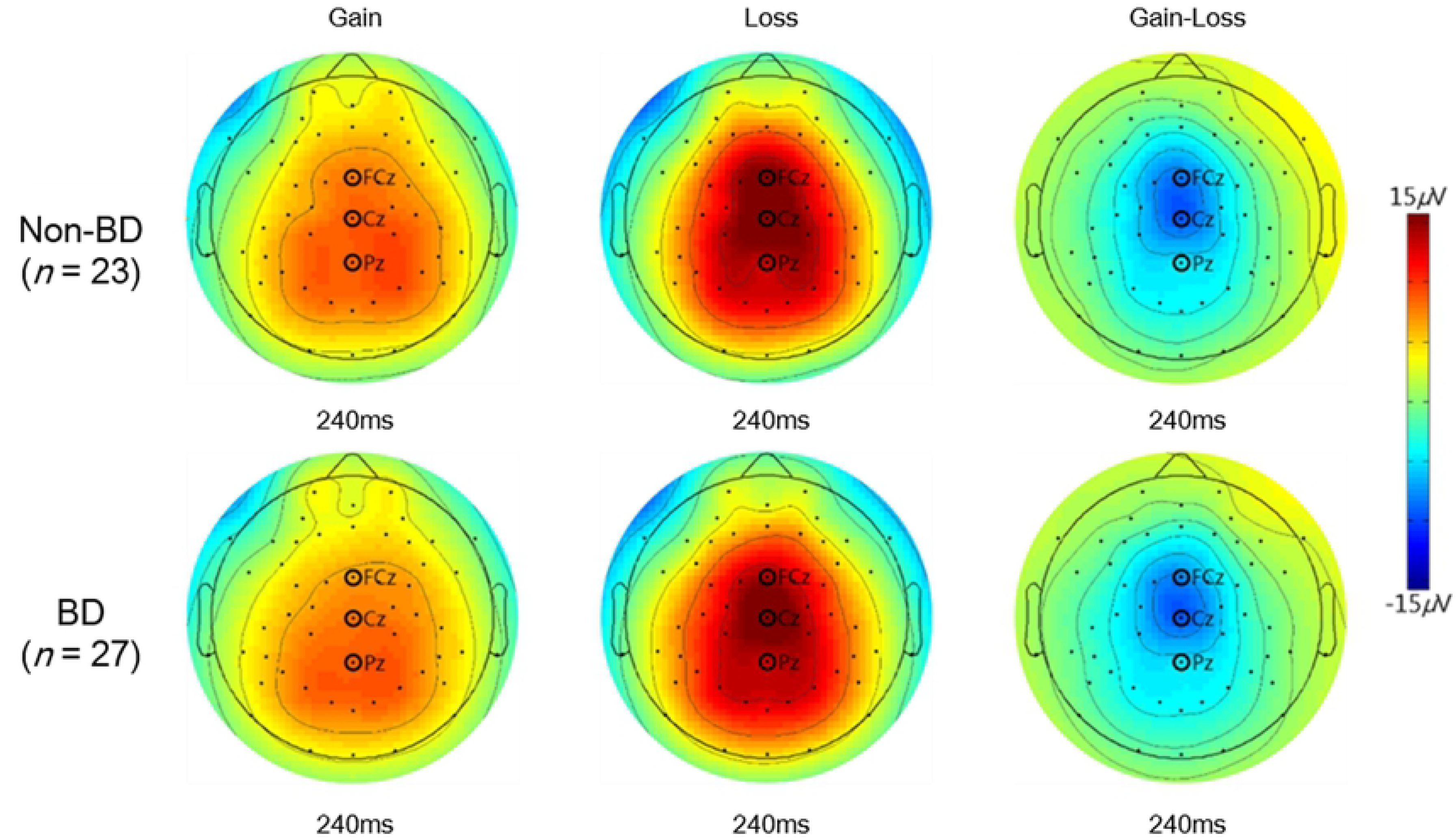
Topographical distribution of P3. The topographical distribution of P3 measured at all electrodes when the maximum P3 amplitudes were observed.

Main effects of valence (F[1,48] = 62.17, *p* < .001) and electrode site (F[8,384] = 18.52, *p* < .001) were observed in terms of FRN amplitudes. FRN amplitudes in response to loss feedback were significantly larger than those in response to gain feedback, and the largest and smallest FRN amplitudes were observed at Cz and FC4, respectively. Additionally, a valence ⅹ group interaction was observed (F[1,48] = 8.06, *p* < .01). A simple effect analysis revealed that while both groups exhibited larger FRN in response to loss than to gain feedback, the difference in the FRN amplitudes between the gain and loss feedback was larger in the non-BD group (mean difference = 2.32, *p* < .001) than in the BD group (mean difference: 1.09, *p* < .01). In addition, an electrode site × valence interaction was observed (F[8,384] = 12.32, *p* < .001) such that FRN amplitudes in response to the loss feedback were larger than those to the gain feedback in all electrodes except FC3. The main effect of group was not significant ((F[1,48] = .08, *p* = .78). The mean FRN amplitudes of the BD and non-BD groups are presented in Table 3.

**Table 3.** Mean FRN amplitudes (*μ*V) in the non-BD and BD groups.

| | Non-BD<br>($n = 23$) | | BD<br>($n = 27$) | |
| --- | --- | --- | --- | --- |
|  | Gain | Loss | Gain | Loss |
| FC3 | 3.41(2.27) | 2.79(2.75) | 3.44(2.02) | 3.43(2.54) |
| FCz | 4.94(3.50) | 1.53(3.84) | 4.51(2.59) | 3.13(3.50) |
| FC4 | 3.32(2.35) | .62(2.56) | 3.38(1.88) | 1.87(2.68) |
| C3 | 3.98(2.60) | 2.80(2.73) | 3.73(1.88) | 3.40(1.97) |
| Cz | 7.11(4.07) | 3.72(4.24) | 6.88(2.65) | 4.67(3.37) |
| C4 | 4.03(2.42) | 1.51(2.48) | 3.48(1.87) | 1.78(2.46) |
| P3 | 4.49(2.50) | 2.96(2.68) | 3.39(1.88) | 3.12(1.82) |
| Pz | 7.04(3.45) | 4.18(3.73) | 6.04(2.65) | 5.33(2.90) |
| P4 | 4.30(2.56) | 1.63(3.11) | 3.50(2.18) | 1.77(3.08) |

**Notes:** () standard deviation.

Main effects of group (F[1,48] = 6.67, *p* < .05) and electrode site (F[8,384] = 12.32, *p* < .001) were observed for ∆FRN. The BD group exhibited a significantly smaller ∆FRN compared to the non-BD group. The greatest ∆FRN amplitude was observed at Cz, and the smallest was detected at FC3. The electrode site × group interaction was not significant (F[8,384] = .70, *p* =.70).

Main effects of valence (F[1,48] = 12.85, *p* < .01) and electrode site (F[8,384] = 3.46, *p* < .05) were observed in terms of FRN latencies. Thus, FRN latencies in response to gain feedback were shorter than those in response to loss feedback (*p* < .01). In addition, the shortest latency was observed at Pz, and the longest was observed at P4. The valence × electrode site interaction was also significant (F[8,384] = 10.25, *p* < .001). The latencies in response to gain feedback were significantly shorter than those in response to the loss feedback at FCz, FC3, FC4, Cz, C3, and C4 but not at the other electrode sites. The valence × group interaction was not significant (F[1,48] = 2.75, *p* = .10). Mean FRN latencies of the BD and non-BD groups are presented in Table 4.

**Table 4.** Mean FRN latencies (ms) in non-BD and BD groups.

| | Non-BD<br>($n = 23$) | | BD<br>($n = 27$) | |
| --- | --- | --- | --- | --- |
|  | Gain | Loss | Gain | Loss |
| FC3 | 218.96(22.23) | 244.17(22.19) | 228.07(28.63) | 235.93(23.10) |
| FCz | 218.70(23.68) | 244.17(17.18) | 229.19(28.19) | 240.07(19.77) |
| FC4 | 226.61(20.22) | 236.70(19.58) | 228.07(21.99) | 238.30(8.53) |
| C3 | 216.96(16.48) | 234.52(23.45) | 227.85(24.95) | 228.89(23.09) |
| Cz | 218.35(20.64) | 238.09(17.71) | 225.41(28.07) | 234.67(17.59) |
| C4 | 228.96(21.11) | 235.39(17.95) | 230.52(19.55) | 234.15(17.51) |
| P3 | 230.26(25.20) | 228.96(24.55) | 221.85(18.68) | 218.59(20.49) |
| Pz | 228.26(23.96) | 220.43(17.44) | 224.15(19.99) | 223.11(21.48) |
| P4 | 236.17(21.23) | 231.48(18.67) | 236.22(16.14) | 230.96(15.77) |

**Notes:** () standard deviation.

Main effects of valence (F[1,384] = 180.72, *p* < .001) and electrode site (F[8,384] = 35.58, *p* < .001) were observed in the P3 amplitudes. The P3 amplitudes in response to loss feedback were larger than those in response to gain feedback, and the largest and smallest P3 amplitudes were observed at Cz and C3, respectively. A valence × electrode site interaction was also observed (F[8,384] = 84.46, *p* < .001), with the largest difference in P3 amplitudes between the gain and loss feedback at Cz and the smallest at P4. However, the main effect of group (F[1,48] = .64, *p* = .43), the interaction effect of valence × group (F[1,48] = .15, *p* = .70), and the electrode site × group interaction (F[8,384] = .59, *p* = .79) were not significant. The mean P3 amplitudes of the BD and non-BD groups are presented in Table 5.

**Table 5.** Mean P3 amplitudes (*μ*V) in the non-BD and BD groups.

| | Non-BD<br>($n = 23$) | | BD<br>($n = 27$) | |
| --- | --- | --- | --- | --- |
|  | Gain | Loss | Gain | Loss |
| FC3 | 5.86(2.90) | 10.49(4.87) | 5.44(2.40) | 10.11(3.46) |
| FCz | 8.4(4.49) | 16.65(7.60) | 6.74(2.90) | 14.63(5.00) |
| FC4 | 7.19(3.47) | 10.92(5.53) | 6.42(2.10) | 9.97(4.16) |
| C3 | 6.02(2.71) | 10.14(4.43) | 5.95(2.25) | 10.05(3.27) |
| Cz | 9.08(4.68) | 19.10(7.69) | 8.18(3.25) | 17.80(5.43) |
| C4 | 7.95(3.42) | 10.75(4.58) | 7.14(2.10) | 10.14(3.58) |
| P3 | 6.80(3.03) | 10.14(3.65) | 6.71(2.22) | 9.98(3.13) |
| Pz | 8.71(4.04) | 13.93(4.22) | 8.96(2.70) | 12.67(4.67) |
| P4 | 8.42(3.43) | 10.62(4.17) | 7.62(2.45) | 10.01(4.04) |

**Notes:** () standard deviation.

A main effect of electrode site was observed in ∆P3 (F[8,384] = 73.33, *p* < .001). The largest ∆P3 amplitude was observed at Cz, and the smallest at P4. No main effect of group (F[1,48] = .23, *p* = .63) or group × electrode site interaction (F[8,384] = 1.08, *p* = .38) was observed.

Main effects of valence (F[1,48] = 51.89, *p* < .001) and electrode site (F[8,384] = 11.65, *p* < .001) were observed for the P3 latencies. The P3 latencies in response to loss feedback were significantly shorter than those in response to gain feedback (*p* < .001); the shortest latency was observed at Pz, and the longest at FC4. An interaction effect of valence × electrode site was also significant (F[8,384] = 7.49, *p* < .001). P3 latencies elicited by loss feedback were shorter than those by gain feedback at all electrode sites except FCz and Cz. The group × valence interaction was not significant (F[1,48] = 2.93, *p* = .09). The mean P3 latencies of the BD and non-BD groups are presented in Table 6.

**Table 6.** Mean P3 latencies (ms) in the non-BD and BD groups.

|  | Non- BD<br>(*n* = 23) |  | BD<br>(*n* = 27) |  |
| --- | --- | --- | --- | --- |
|  | Gain | Loss | Gain | Loss |
| FC3 | 336.70(35.14) | 324.17(22.20) | 334.89(28.16) | 315.93(23.10) |
| FCz | 329.57(32.75) | 324.17(17.18) | 328.00(30.47) | 320.07(19.77) |
| FC4 | 334.70(31.24) | 316.70(19.58) | 342.30(25.59) | 318.30(18.53) |
| C3 | 339.39(35.19) | 314.52(23.45) | 346.30(30.74) | 308.89(23.09) |
| Cz | 317.83(32.79) | 318.09(17.71) | 324.30(36.81) | 314.67(17.59) |
| C4 | 336.43(29.55) | 315.39(17.95) | 346.37(24.51) | 314.15(17.51) |
| P3 | 334.26(33.27) | 308.96(24.55) | 337.26(33.62) | 298.59(20.49) |
| Pz | 311.13(29.54) | 300.43(17.44) | 316.37(35.65) | 303.11(21.48) |
| P4 | 329.48(29.67) | 311.48(18.67) | 348.74(27.94) | 310.96(15.77) |

**Notes:** () standard deviation.

### Correlations between performance on the modified IGT and ∆FRN/∆P3 amplitudes

Positive correlations were observed between ∆FRN amplitudes at FCz and total net scores (r = .298, *p* < .05), sector 4 net scores (r = .333, *p* < .05), and sector 5 net scores (r = .357, *p* < .05) of the IGT. Thus, larger ∆FRN amplitudes at FCz were associated with better IGT performance, especially in the later sectors of the IGT. On the other hand, no significant association was detected between the ∆P3 amplitudes and IGT performance.

## Discussion

This study investigated feedback utilization ability for decision making in BD college students using the modified IGT and ERP data. The BD group exhibited significantly lower total net IGT scores and lower ∆FRN amplitudes than did the non-BD group. Additionally, the ∆FRN amplitude at the fronto-central area was positively correlated with the total net scores, sector 4 net scores, and sector 5 net scores on the IGT.

The BD group exhibited significantly lower total net scores than the non-BD group did, and performance of the non-BD group tended to increase as the task progressed (mean sector 1 = -.03; sector 2 = .73; sector 3 = .93; Sector 4 = 2.23; sector 5 = 1.25), whereas the BD group persistently chose disadvantageous cards over advantageous ones (mean sector 1 = −1.73; sector 2 = −3.04; sector 3 = −1.63; sector 4 = −1.41; sector 5 = −1.93). These results are consistent with those of previous studies [30–33] and suggest that individuals with BD have deficits in decision making. To maximize gains on the IGT, one must choose more advantageous cards that provide small initial gains but result in a net gain over disadvantageous cards that provide a large initial gain but result in a net loss. Johnson et al. [30] suggested that poor performance on the IGT in individuals with BD reflects their failure to consider consequences, i.e., tendency to pursue immediate rewards, disregarding the larger potential risk.

The statistical analyses of FRN, one of the ERP components elicited by feedback, revealed that the BD group exhibited significantly lower ∆FRN amplitudes than the non-BD group did. The non-BD group exhibited larger FRN amplitudes in response to loss feedback than to gain feedback, whereas the FRN amplitude differences in the BD group between gain and loss feedback were significantly smaller than those in the non-BD group. These results are consistent with previous studies on patients with AUD and male BD college students [69–71]. The present study also revealed that both groups exhibited larger FRN amplitudes in response to loss feedback than to gain feedback, which is consistent with many previous studies [58,60,97-99] and suggests that FRN is sensitive to feedback valence. FRN is known to reflect an early evaluation of feedback provided by the environment [48,60,98]. For example, Yeung and Sanfey [60] suggested that FRN and P3 reflect early and late stages of feedback processing, respectively. Gu et al. [98] reported that FRN reflects early feedback evaluation based on the salience of the feedback information.

Insensitivity to future consequences (IFC) in patients with AUD and substance use disorder (SUD) has been consistently reported [21,100,101]. For example, Cantrell et al. [101], using a modified version of the IGT, measured the preference for larger versus smaller rewards (PLvS), the difference between frequencies of choosing the cards that provide large gains and cards with small gains, and IFC, the difference between frequencies of choosing cards that result in a net loss and cards that result in a net gain in patients with AUD. The results showed that although patients with AUD did not exhibit significantly different PLvS scores, they exhibited significantly higher IFC scores than the control group. Additionally, a study of patients with SUD, including AUD, using the IGT and the Prospect Valence Model analysis observed a consistent lack of sensitivity to losses in patients with SUD [102]. Therefore, significantly smaller ∆FRN amplitudes in the BD group compared to the non-BD group observed in the present study suggest that the BD group has deficits in early feedback evaluation and that they are less sensitive to loss feedback than are members of the non-BD group.

In this study, no significant difference in the P3 amplitudes was observed between the BD and non-BD groups, which was not consistent with previous studies reporting reduced P3 amplitudes in patients with AUD [70,103]. The generators of P3 are known to be located in the temporo-parietal junction or locus coeruleus-norepinephrine system [67]. On the other hand, alcohol is known to affect frontal areas of the brain [68,104]. For example, those who consume alcohol exhibit reduced N450 amplitudes in frontal areas, whereas P3 amplitudes in the parietal and occipital areas are not affected by alcohol consumption [104]. Nelson et al. [68] also reported that alcohol consumption reduces both theta and delta band activities, which are known major components of FRN and P3, respectively, affecting theta band activity more severely. Whereas alcohol consumption affects the frontal area, overall grey and white matter volume reductions, including those in frontal areas, are observed in patients with AUD [46,105,106]. For example, one study observed reduced whole-brain network cluster coefficients in patients with AUD and reported that longer AUD duration was associated with a global decrease in the efficiency of the brain network [107]. These results suggest that alcohol consumption affects frontal areas first, and then spreads over the whole area as drinking duration increases. Taking together, our results imply that BD of relatively short duration (the mean drinking duration in the BD group was 33.33 months) may affect later feedback evaluation and attentional resource allocation relatively less severely than does BD with a long drinking history.

Both groups exhibited larger P3 amplitudes with loss feedback than with gain feedback. Studies on feedback-related ERPs using tasks other than the IGT have reported larger P3 amplitudes in response to gain feedback than to loss feedback [63,64,108,109], whereas studies using the IGT observed larger P3 amplitudes in response to loss feedback than to gain feedback [93,110]. Feedback-related P3 is known to be sensitive to different feedback information, not just to feedback valence but also to feedback magnitude and probabilities as well [60,63,64,66,97,98,111]. This suggests that P3 reflects feedback processing with a top-down mechanism that allocates attentional resources to the information relevant to the task at hand [67,98]. The loss magnitudes of each card must be understood to maximize profit on the IGT. Thus, participants need to understand that disadvantageous cards (A and B) result in large gains, but losses will soon accumulate over gains, and thus shift their preference or attention progressively toward advantageous cards (C and D) [44]. These results suggest that both groups allocated their attentional resources to feedback valence, especially to loss feedback, while taking the modified IGT.

Although the importance of feedback utilization for decision making has been emphasized [34,48], only one study has investigated the association between IGT performance and feedback-related ERPs [93]. Carslon et al. [93] investigated how children responded to gain/loss feedback using P3 and evaluated how anticipation prior to the response was related to behavioral adjustment using stimulus-preceding negativity (SPN). That study found that the difference in children’s SPN amplitude between advantageous and disadvantageous decks was positively correlated with behavioral adjustment. In the present study, ∆FRN amplitudes at FCz were positively correlated with total net scores and sectors 4 and 5 net scores on the IGT. Thus, larger differences between FRN amplitudes with gain and loss feedback were associated with improved performance on the modified IGT. No previous study has reported an association between FRN and IGT performance; studies using the reversal learning task have reported associations between FRN and behavioral adjustments [108,112]. For example, Frank et al. [112] compared negative learners, who learn stimulus-result contingencies by avoiding negative feedback, with positive learners, who learn these contingencies by pursuing positive feedback; they found significant positive correlations between the tendency to avoid negative feedback and error-related negativity (ERN) amplitudes, which is the ERP component known to share some neural sources with FRN [94]. To perform successfully on the IGT, participants must learn the contingencies between the cards and their consequences implicitly during the task (e.g., gain or loss feedback) [113]. Therefore, these results suggest that early feedback evaluation in the fronto-central area is associated with the implicit learning process during decision making.

No significant associations between ∆P3 amplitudes and IGT performance were observed in this study. Previous results for P3 amplitudes and behavioral adjustments using the reversal learning task are inconsistent [112,114]. For example, Frank et al. [112] reported that FRN amplitudes, not P3 amplitudes, predicted behavioral adjustment, whereas Chase et al. [114] reported that P3 amplitudes, not FRN amplitudes, predicted behavioral adjustment. The differences between these two studies lay in the task instructions. Frank et al. [112] did not provide any information regarding contingency shifts during the task and requested that participants make decisions based on their internal judgment, whereas Chase et al. [114] told the participants that the contingency would shift during the task and requested that participants adjust their responses when they were certain that the contingency had shifted. Thus, the latter study reflected decision making based more on a set of rules provided prior to the task than on the actual feedback during the task. San Martin [48] suggested that the importance of FRN and P3 in behavioral adjustment varies depending on which information is more important when performing the given task. In our study, participants were only instructed regarding the goal and process of the task. Therefore, the rules of the task (probability of loss and magnitude of cards) must be learned solely through feedbacks. Such a task design is closer to the study by Frank et al. [112]. These results suggest that both the BD and non-BD groups relied more on early feedback evaluation of valence than on late evaluation with a top-down mechanism as they performed the modified IGT.

This study has several limitations. First, the feedback evaluation investigated here focused mainly on feedback valence. The magnitudes of the cards on the modified IGT increased, as was the case in the original IGT [41]. Although this may keep participants motivated, the increasing magnitude forbids examination of how feedback-related ERPs respond differently to feedback of small or large magnitude. Additionally, the probabilities of encountering losses from cards B and D were too low (10%) to secure enough trials to investigate how ERPs differ based on the probability of losses. Second, this study measured feedback utilization using time-based ERPs. However, the time windows for FRN and P3 are close to each other, and they may distort each other in ERP waveforms. Difference waves were measured to isolate ERP components and prevent such distortion, but other techniques, such as spectral analysis or functional connectivity analysis, may reveal more detailed information, such as how different neural waves interact and communicate during feedback processing.

In conclusion, the BD group exhibited significantly lower total net scores on the modified IGT and significantly lower ∆FRN amplitudes. On the other hand, no differences were observed in ∆P3 or P3 amplitudes between the groups. Additionally, positive correlations were observed between ∆FRN amplitudes in the fronto-central area and IGT performance. These results imply that the BD group had deficits in decision making and early feedback evaluation, with a tendency to pursue immediate large gains even at greater potential risks, revealing deficits in early evaluation regarding feedback valence.

## Author Contributions

**Conceptualization:** Myung-Sun Kim.

**Formal analysis:** Eun-Chan Na, Kyoung-Mi Jang.

**Funding acquisition:** Myung-Sun Kim.

**Methodology:** Eun-Chan Na, Kyoung-Mi Jang.

**Supervision:** Myung-Sun Kim.

**Writing – review & editing:** Eun-Chan Na, Kyoung-Mi Jang, Myung-Sun Kim.

## Supporting information

**S1 File. Behavioral data.**

(XLSX)

**S2 File. ERP data.**

(XLSX)

